# Integration of heterogeneous biological data in multiscale mechanistic model calibration: application to lung adenocarcinoma

**DOI:** 10.1101/2022.01.17.476676

**Authors:** Jean-Louis Palgen, Angélique Perrillat-Mercerot, Nicoletta Ceres, Emmanuel Peyronnet, Matthieu Coudron, Eliott Tixier, Ben M.W. Illigens, Jim Bosley, Adèle L’Hostis, Claudio Monteiro

## Abstract

Mechanistic models are built using knowledge as the primary information source, with well-established biological and physical laws determining the causal relationships within the model. Once the causal structure of the model is determined, parameters must be defined in order to accurately reproduce relevant data. Determining parameters and their values is particularly challenging in the case of models of pathophysiology, for which data for calibration is sparse. Multiple data sources might be required, and data may not be in a uniform or desirable format. We describe a calibration strategy to address the challenges of scarcity and heterogeneity of calibration data. Our strategy focuses on parameters whose initial values cannot be easily derived from the literature, and our goal is to determine the values of these parameters via calibration with constraints set by relevant data. When combined with a covariance matrix adaptation evolution strategy (CMA-ES), this step-by-step approach can be applied to a wide range of biological models. We describe a stepwise, integrative and iterative approach to multiscale mechanistic model calibration, and provide an example of calibrating a pathophysiological lung adenocarcinoma model. Using the approach described here we illustrate the successful calibration of a complex knowledge-based mechanistic model using only the limited heterogeneous datasets publicly available in the literature.

## 1 Introduction

Multiscale mechanistic modeling is an approach that bridges physiology and pathophysiology across multiple spatial scales and physical processes (Eissing, 2011; Buil-Bruna et al, 2015; Musante et al, 2016; Boissel et al, 2015) and is also called knowledge-based modeling. Knowledge-based models (KBMs) capture the complexity of biological functions in one or several subparts of a mechanistic model, using a broad range of knowledge (see Box 1) extracted from data.

### Box 1: Definitions of modeling-related terms in dynamic biological models with only time dependency

#### Parameter

quantitative entity within a model whose value is set to remain constant in a particular context. For example, an IC50 value might represent the drug concentration yielding half-maximal tumor growth inhibition.

#### State variable

quantitative entity within a model, which value depends on parameters and/or other variables and is expected to vary with time. For instance, the number of cells in a tumor may vary in response to treatment or immune system activity. For the sake of simplicity, state variables are referred to as variables in the remainder of the article.

#### Knowledge

structured information about a phenomenon that is derived from several observations and/or experiments. It can refer to an entity (*e*.*g*. an ionic channel, a gene), a relationship (*e*.*g*. an ionic current which brings a signal from entity X to entity Y) or a phenotype/genotype for instance. A piece of established knowledge, also called a scientific fact (*e*.*g*. the immune system consists of several cell types) is based on multiple observations and experiments (*i*.*e*. data) and is therefore considered at the top level of evidence.

#### Virtual Patient

vector of parameter values that can characterize an individual from the model perspective.

#### Phenomenon

biological process defined by its given inputs, outputs, and granularity level (*e*.*g*. a metabolic pathway or circulation of one entity).

#### Data

structured observation with qualitative or quantitative information from a defined set of subjects (*e*.*g*. patients). It refers to the information obtained from a single statistical unit in an observational study (*e*.*g*. cohort, survey, -omics study) or an experiment (*e*.*g*. animal experiment, *in vitro* experiment, clinical trial).

KBMs rely on mathematical equations composed of parameters to mechanistically represent biological phenomena through the changes over time of biological variables (Box 1). Many of these variables can be measured experimentally (*e*.*g*. evolution of glucose concentration in the plasma), while others are difficult to quantify in an experimental settings (*e*.*g*. concentration of intracellular ATP in a specific cell) (Brown et al, 2018). To confidently draw conclusions from modeling analysis, the measurable variables of the model must reproduce what is observed in experimental settings, and the knowledge used to link measurable and non-measurable variables must be consistent with the scientific consensus. While the latter is ensured if the model is well constructed, the former can only be achieved by defining and adjusting parameter values that enable model measurable variables to reproduce the desired behavior, that is to say what is observed in experimental settings. Two methods are commonly used to find the parameter values (Box 1): parameterization and calibration (Box 2).

### Box 2: Definitions of calibration-related terms

#### Parameterization

selection of parameter values based on human literature curation, either directly referenced values or extrapolation using assumptions (*e*.*g*. taking a similar value from other analogous biological entities).

#### Biological constraint

biological behavior identified in experimental data or knowledge, that the model is designed to reproduce.

#### Computational constraint

computational translation of a biological constraint into numerical expression. Each computational constraint can be evaluated in a given virtual patient and generates an associated score (see calibration score).

#### Calibration

an automatic process in which parameters values that are *a priori* unknown are refined through numerical optimization routines run in parallel in order to achieve a desired behavior of the model.

#### Calibration score

when a computational constraint is evaluated, a numerical value or score between 0 (the constraint is not fulfilled) and 1 (the constraint is fully fulfilled) is generated for each virtual patient. Each score can be weighted to reflect a user-defined relative importance compared to other scores. For a given virtual patient, the objective function to be maximized is expressed as the weighted sum of all its scores.

#### Parameter space

number of parameters and their possible set of values within the model. Because parameters are based upon biology, their set of possible values is limited to what is biologically possible. The size and shape of the parameter space thus refers to both the number of parameters, their possible values and the complexity of their interactions (*e*.*g*. linear vs non-linear interactions).

The term parameterization can be applied to describe human curation used to search the scientific literature and to select representative parameter values. In contrast, calibration is an automatic numerical procedure, in which *a priori* unknown and estimated parameter values are concomitantly refined to reproduce the desired model’s behavior, usually via an iterative process. Calibration constrains the dynamic behavior of the model (*e*.*g*. during simulation) by finding a set of parameter values that allows the model to represent biological behaviors consistent with the literature. On one hand, parameterization allows for an objective assignment of a parameter value avoiding overfitting of a dataset, while one step calibration could bring new information to fix parameters with no reported value. On the other hand while parameterization can be appropriate for models with a small number of parameters, calibration allows one to address models featuring a large number of parameters, that may have strong interactions (Box 2). Multiscale mechanistic KBMs are generally part of this second category of models and thus are subject to calibration (Box 3).

### Box 3: Definitions of modeling approaches and model constraints

#### Knowledge-based models (KBMs) vs data-driven models (DDMs)

The difference between KBMs and DDMs lies in how the models are founded/implemented: mathematical equations that represent knowledge on each biological interaction of interest for the former, and a statistical model that is assumed to fit with the data and the way they were collected for the latter. As such, KBMs are composed of parameters and variables that have plausible physiological meaning.

#### Calibration *vs* fitting

Model fitting, as used in DDMs, is often based upon unconstrained modification of both the model structure and the related parameters to minimize a metric of fit representing the difference between measured biological entities and their modeled equivalent. By contrast, calibration is more adapted to KBMs composed of parameters and variables that have physiological meaning. It is applied on a model structure deduced from knowledge, and consists in minimizing the difference between measured physiological data and modeled variable dynamics upon constrained parameter value modification, based on variable behavior. As a consequence, while fitting is usually performed on a limited number of variables (e.g. drug concentration and clinical outcome) and can change the model structure, calibration explores an additional set of constraints (such as model variables remaining physiologically plausible) to ensure more robust predictions, keep the model structure and is able to filter variable acceptable behavior.

The objective of KBM calibration is to find the set of parameter values that allow the model variables to reproduce the desired biological behavior, that is to say a set of qualitative and quantitative knowledge and data extracted from the literature. However, knowledge and data found in literature are scarce, therefore the aim of this article is to establish a calibration strategy for existing non-calibrated computational models, using publicly available heterogeneous data to accurately estimate model parameters’ values or distributions. Heterogeneity in this context includes several aspects of the data, including both the different scales of the data (some obtained *in vitro*, some in animal model, some in humans), different types of outputs (distinct variables measures) and whether the measurements are qualitative or quantitative. During the calibration process, the model architecture itself is assumed to remain static (*i*.*e*. no refactoring of the model, such as adding, removing or replacing a biological phenomenon is allowed to occur), and only the parameter values are changed. In this paper, we present a calibration methodology where the benefits of heterogeneous types of data are leveraged through (1) data assessment, (2) assurance of satisfying adequacy between model and observed data, (3) definition of the aim of each calibration step, (4) automatic optimization of parameters, and (5) integration of all calibration steps. The resulting calibration protocol addresses (1) data with different degrees of precision, from individual measurements to general knowledge, (2) data with different provenance (*in vitro* or *in vivo* and from several species), (3) data with different granularity (individual record or population-based), and (4) calibration at different levels, *i*.*e*. calibration of all measurable variables, regardless of their granularity.

This general calibration strategy is then applied to a multiscale mechanistic model of lung adenocarcinoma (LUAD). This study focuses only on the calibration application and will not address how model predictivity can be assessed on external data for a validation purpose.

## 2 Methods

### 2.1 Mechanistic modeling

Mechanistic models are based on knowledge retrieved from literature, and represent, notably through model parameters values, well-described biological relationships between all model variables, the so-called biological phenomena (Box 1). The values of the model parameters therefore need to be correctly estimated for the model to reproduce the biology. The initial parameter identification provides plausibility of value ranges for parameters and should as much as possible be assessed on physiological knowledge. Initial values can be generated or inferred using known relationships or by hypotheses (*e*.*g*. via analogy or scaling). Parameters have been classified in distinct categories in literature based on the way their initial value is estimated (Yugi, 2013). Here we propose a classification in two classes according to whether the parameter value is set through parameterization or calibration.

In this article, we will focus parameters that require calibration to estimate their values to reproduce desired biological behaviors. Finding the data required to represent such expected biological behaviors requires a thorough review of the scientific literature. This step is nevertheless needed to obtain qualitative and quantitative knowledge and data that can be used to constrain the model behavior. A model is said to be calibrated when the computational constraints derived from the review are achieved.

The information found in literature usually follows one of these three formats: text descriptions, numerical values, or graphical values. All three can be converted into computational constraints. This step is crucial, as the predictive power of a mechanistic model depends on the quality and consistency of the computational constraints on which the model is evaluated. The exercise is also critical in ensuring that models fulfill biological constraints that are meaningful to the future application of the model.

Prior to the calibration *per se*, it is key to identify which parameters to calibrate, trying to reduce their number at most to ease the upcoming calibration, as well realistic ranges for their values. This can sometimes be done during literature search by defining extreme values the parameter cannot take. Most often this can be done using sensitivity analysis methods, many of which can be applied in multiscale models (Renardy et al, 2019). Here, we use regular factorial designs to perform a multivariate sensitivity analysis (Monod et al, 2012; Bidot et al, 2018). Briefly, it allowed us to explore large ranges of parameter values, discard those that led to either numerical error or aberrant model behavior (defined based on literature review) and thus define the parameter space to explore.

### 2.2 Integration of heterogeneous data with a step-by-step calibration

The adequacy between the model variables and the relevant biological entities is a matter of ensuring that the model operates within the computational constraints and reproduces the expected behaviors. Depending on the level of knowledge available on some phenomena, distinct scales or levels of details (*e*.*g*. molecular or cellular) may coexist in a single model, notably in multiscale models. Multiscale models describe a complex process across multiple scales of time and space, and account for how quantities transform as we move from one scale to another (Bhattacharya and Viceconti, 2017; Horstemeyer, 2009).

The knowledge and data used to define the computational constraints are heterogeneous and can have different levels of granularity. To overcome this discrepancy, it is then relevant to approach the calibration of a model as a list of successive steps, each step having as its objective a specific model variable behavior matching one or more specific computational constraints. Since calibration steps are executed sequentially, the first calibration steps are prerequisites for the following steps. The order of the calibration steps is therefore important and the first steps should focus on reproducing the computational constraints related to:

- biological phenomena that are well-detailed in the literature, in order to introduce as few uncertainty as possible in the first calibration steps (as they would have repercussions on the following steps)
- biological phenomena represented with the lowest level of granularity to ensure that the most detailed variables, and often the most sensitive to parameter value changes, have a consistent behavior in the model
- biological phenomena that are less connected to the others, such as a subpart of the model that is less connected to the others to ensure that a calibration solution within the parameter space can more easily be found (compared to the exploration of a wide parameter space with deeply interconnected models). In this context, connection refers to the number of inputs/outputs between the models: the less inputs from other models a given model needs to run, the less connected it is considered

Most often, at each step one calibrates subparts of the model (also called submodels). Later steps combine these submodels to calibrate larger and larger models.

Each of the calibration steps reduces the remaining space of the parameters to be estimated. A calibration step starts with a number of parameters for which we have large uncertainty (a wide range of possible values). The calibration then either sets a definite value for that parameter (reducing the number of unknowns in the consecutive calibration steps) or it reduces the possible range of values for a parameter, limiting the plausible parameter space. Some computational constraints may influence multiple calibration steps, as each new step in the calibration should ensure that the results from the previous steps are still relevant; *i*.*e*. the constraints used at one step should still be verified in subsequent steps. Overall, the feasibility of a model calibration depends on the amount and quality of the experimental data used to define the computational constraints. An illustration of such a calibration strategy is provided in Figure 1.

**Fig. 1.**
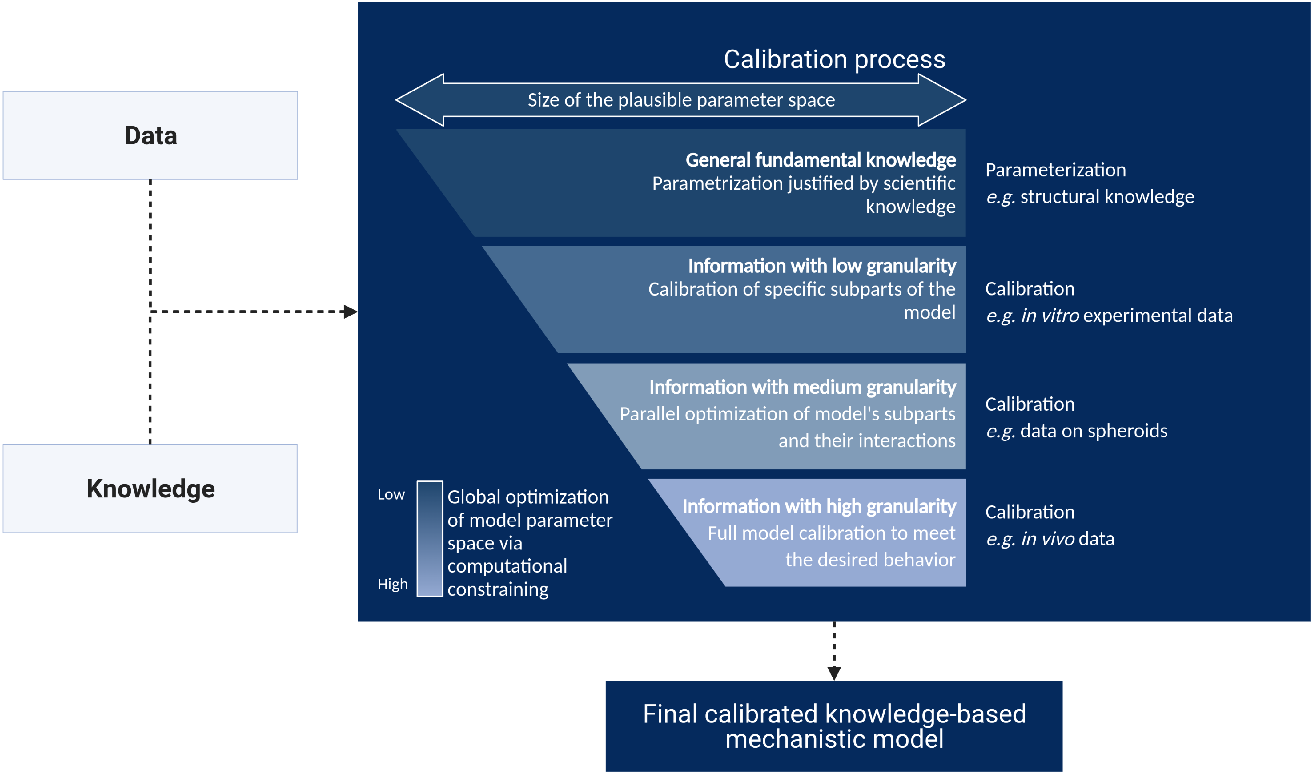
Illustration of the stepwise calibration process of a model with knowledge and data as input. We propose an iterative calibration process where the size of the plausible parameter space decreases at each step, and the effect of the computational constraints applied to the model on the efficiency of the calibration process increases.

At the end of a calibration step, the model parameter values should ensure that the model reproduces the biological behavior defined by the computational constraints.

At the end of the calibration process the model should be able to reproduce the pathophysiology of interest, in the desired context of use, represented by the whole set of computational constraints (Morrison et al, 2019).

### 2.3 Computational implementation of calibration

#### 2.3.1 Overview of existing methods

Calibration, or parameter estimation, consists in solving the inverse problem of finding parameters values such that the model matches the available experimental data (Moles et al, 2003). Most calibration methods are variations of either a Bayesian inference or a maximum likelihood problem (Klipp et al, 2016). The principle of Bayesian inference is to estimate the probability distribution of the parameters (the posterior distribution) given the probability of observing the available data (the likelihood) while taking into account prior belief (the prior distribution) regarding the parameters distributions (Liepe et al, 2013). Bayesian inference is particularly suited to problems involving scarce, missing or noisy data however estimating the posterior distribution is often analytically intractable, therefore sampling methods such as Monte-Carlo Markov Chains (Rosenthal, 1995) can be used. Even then, the computational cost can remain prohibitive in high-dimensional cases which is why methods such as Approximate Bayesian Computation (Barber et al, 2015; Toni et al, 2009) or Sequential Monte Carlo (Liu and Chen, 1998) have been developed to provide less costly likelihood approximations. Maximum likelihood methods aim at estimating the parameters that maximize the goodness of fit of a model, *i*.*e*. minimize the discrepancy between model outputs and experimental data. They usually amount to maximizing a non-linear objective function for which there exists many algorithms that can be divided in two classes: deterministic and stochastic. Deterministic methods can involve the gradient (Galtier and Wainrib, 2013) of the objective function which makes them more likely to converge to local maxima. They can also be “gradient-free” such as the DIRECT Jones et al (2001) and Nelder and Mead (1965) methods for instance. Stochastic methods generally tend to be more robust with respect to local maxima while being relatively easier to implement when they treat the objective function to maximize as a black box. Some examples are simulated annealing (Kirkpatrick et al, 1983), the SAEM algorithm (Delyon et al, 1999), which is particularly suited to non-linear mixed effects models and evolutionary methods which have the advantage of being easily parallelizable (Back et al, 1997).

#### 2.3.2 Implementation of calibration with CMA-ES

CMA-ES, which stands for Covariance Matrix Adaptation Evolution Strategy (Hansen et al, 2003; Jdrzejewski-Szmek et al, 2018; Tomasoni et al, 2021) is a member of the aforementioned evolutionary methods which has shown to be efficient for a wide variety of applications (Ryan et al, 2007).

Akin to most evolutionary methods, CMA-ES proceeds iteratively, each iteration corresponding to a generation of *p* individuals, each individual being a vector of the parameter values to optimize. For a given generation, the objective function is evaluated for each individual, a subset of individuals exhibiting the best objective values is kept (it is the “survival of the fittest” principle which gave evolutionary methods their name) and is used through reproduction to generate the next generation. In the CMA-ES algorithm, the individuals of a generation are obtained by sampling from a multivariate normal distribution and the reproduction step consists in using the fittest individuals to update the covariance matrix and mean of said distribution. The hyperparameters of the algorithm are *p*, the number of individuals per iteration, the initial covariance matrix and mean and the stopping criterion.

In the context of this article, the goal of calibration is to obtain parameter values such that the computational model satisfies some biological constraints. As shown in Figure 2, constraints can be of two kinds: a knowledge source (e.g. a quantity of interest is above a given threshold) with an associated binary score that takes value 0 or 1 or a data source (*e*.*g*. a measure of how close an output variable dynamics is with respect to experimental data) with a score which continually varies in [0, 1]. The motivation for representing constraints with such scores is to have quantities of comparable orders of magnitude such that they can seamlessly be combined by weighting their sum into a single objective function which itself varies in [0, 1]. Rescaling the scores and objectives in [0, 1] also allows for easy interpretation of the result.

**Fig. 2.**
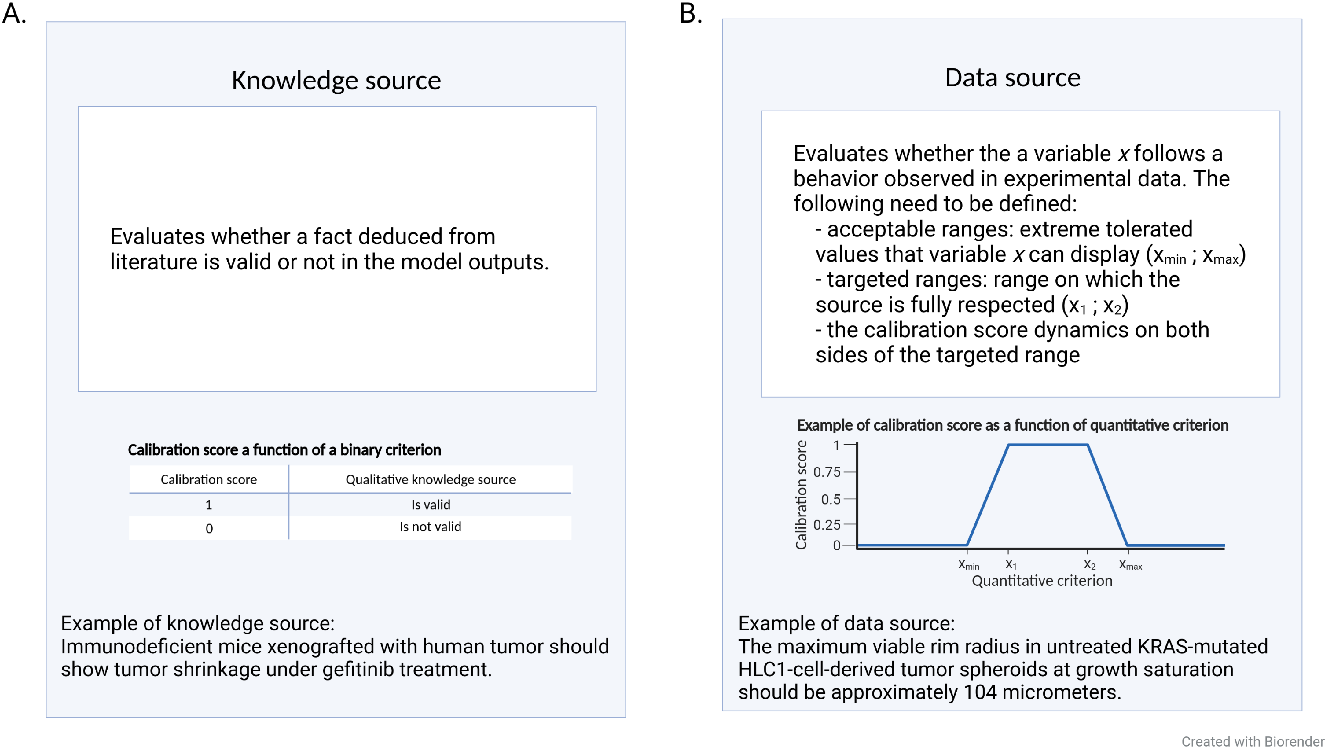
Translation of biological constraints into computational constraints. A. Computational constraint based on knowledge, B. Computational constraint based on experimental data. Reference (Freyer, 1988).

Each computational constraint is weighted to reflect a user-defined importance compared to the others. For a given virtual patient, the objective function *f* (*θ*) to maximize is the weighted sum of all its scores and reads:

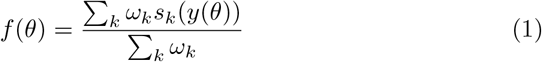

where *y*(*θ*) is the simulation output, *s*_*k*_ the k^th^ sub-score and *ω*_*k*_ its associated weight.

The objective function f(*θ*) is maximized using the CMA-ES algorithm. The population size hyperparameter *p* is chosen using the rule recommended by the Python library implementing CMA-ES (Hansen et al, 2019):

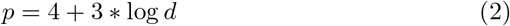

where *d* is the number of parameters to calibrate

The initial mean of the multivariate normal distribution corresponds to the individual initial guess of the parameters which are user-defined. The initial covariance matrix is a diagonal matrix where the square root of the k-th diagonal coefficient (which corresponds to a standard deviation) is chosen such that it equals half of the k-th parameter search interval width.

The stopping criterion is whatever is satisfied first among the following two:

- the objective function reaches a user-defined threshold (typically close to 1)
- the objective function stagnates over multiple iterations (Hansen and Kern, 2004)

Calibration is deemed satisfactory when the model outputs accurately reproduce the biological behavior depicted by the computational constraints (Allen et al, 2016).

#### 2.3.3 Illustration with a synthetic example

In this section we propose to illustrate the calibration methodology presented in the above with a toy example consisting of a cost function:

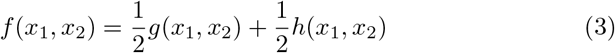

where

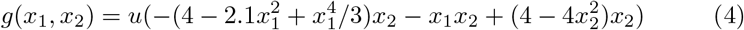

and

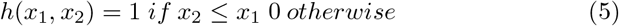

*f* is a simple example of objective function combining one binary and one continuous score with equal weights. *u* is a rescaling function such that *g*(*x*_1_, *x*_2_) takes values in [0, 1]. *g*(*x*_1_, *x*_2_) has a known global maximum of 1, reached twice in (*x*_1_, *x*_2_) = (± 0.0898,∓ 0.7126) and four local maxima.

The proposed calibration procedure is applied to this toy example with initial guess (*x*_1_, *x*_2_) = (−2, 1), initial standard deviations *σ*_1_ = *σ*_2_ = 0.5 and population size *p* = 20.

Figure 3 illustrates how the calibration problem is solved using CMA-ES. The samples converge towards the global maximum which corresponds to one of the global maxima of *g*(*x*_1_,_*x*_ 2) where *h*(*x*_1_, *x*_2_) = 1.

**Fig. 3.**
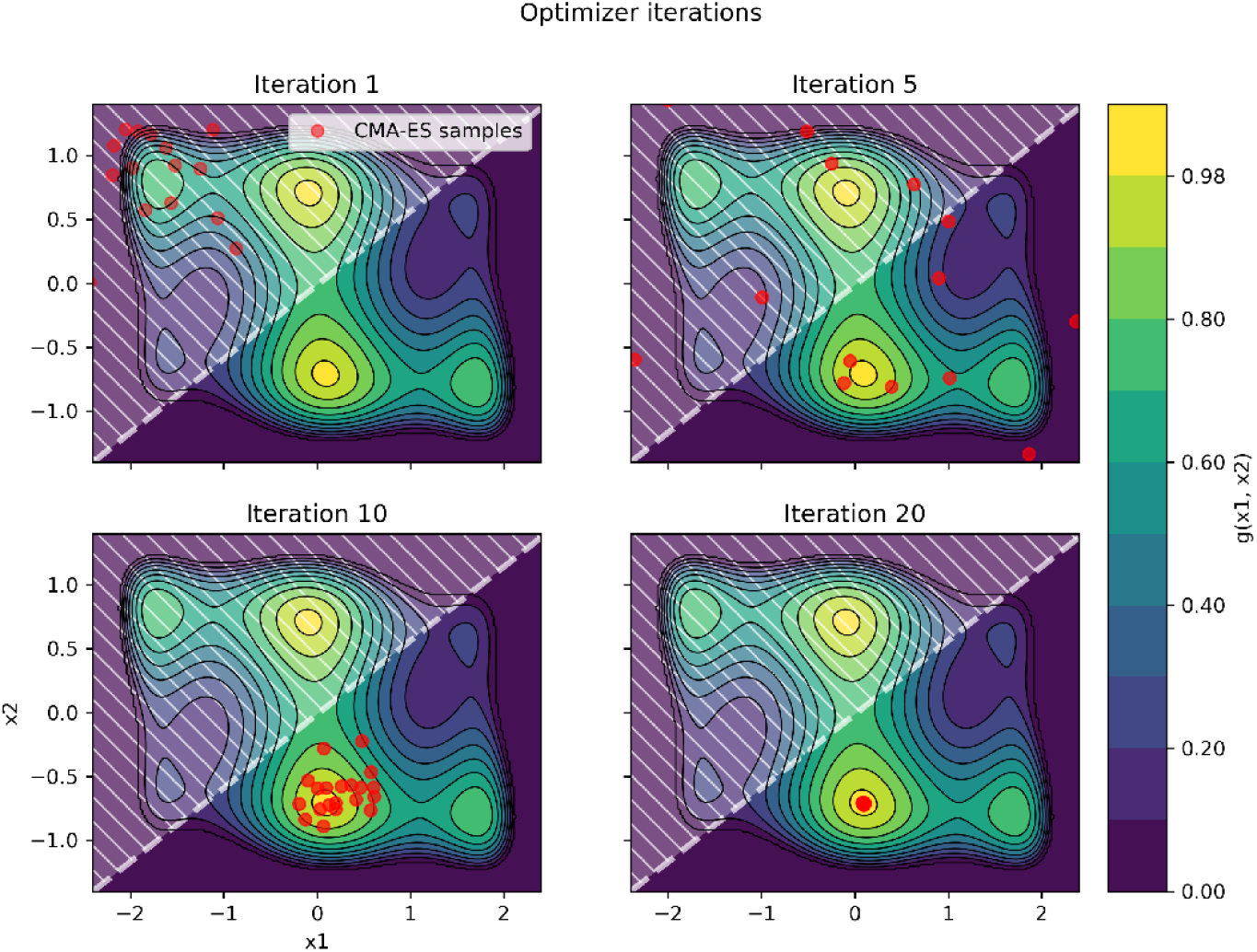
Illustration of the convergence of CMA-ES on a toy calibration example. Contours correspond to the continuous score *g*(*x*_1_, *x*_2_)and the dashed area materializes where the binary score *h*(*x*_1_, *x*_2_)equals zero. The red dots show the CMA-ES samples at different iterations.

Figure 4 shows the evolution of the objective function values associated with the CMA-ES samples throughout iterations.

**Fig. 4.**
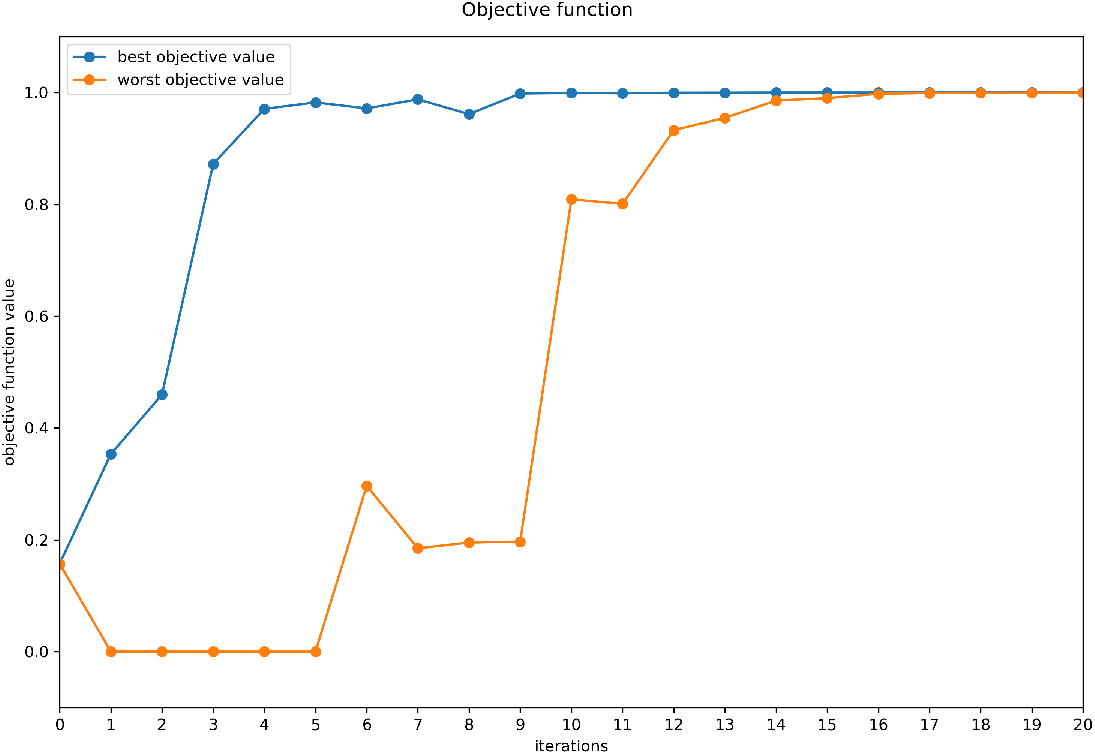
Evolution of the objective function with CMA-ES iterations. For each iteration, the CMA-ES algorithm draws and evaluates the score (or objective value) of several sets of parameter values, sampled around the best set of parameter values from the previous iteration. The worst and best samples for each iteration are plotted to illustrate that a) CMA-ES finds in few iterations regions with high objective values and b) the drawn samples converge to the neighborhood of the global maximum.

### 2.4 Parameter identifiability

The present paper does not deal with parameter identifiability. The aim is not to find a unique solution to the calibration problem but rather find one set of parameters among several which are assumed all likely to give a good fit. It can even be argued that even if the estimation process is not able to tightly constrain any of the parameter values, the model can still be able to yield significant quantitative predictions (Brown and Sethna, 2003). The interested reader is referred to Duchesne et al (2019), Miao et al (2011) and Pant and Lombardi (2015) for studies regarding parameter identifiability in dynamical systems.

## 3 Application example

### 3.1 Context: mechanistic model of lung adenocarcinoma under gefitinib treatment

Non-small-cell lung cancer (NSCLC) is the most common type of lung cancer accounting for 80-90% of all lung cancers, and about 40% of all lung cancers are adenocarcinomas (LUAD)(ESMO, 2019). These tumors start in mucus-producing cells that line the airways. The studied model, referred to as the *in silico* EGFR+ (Epidermal Growth Factor Receptor) Lung Adenocarcinoma (ISELA) model, relies on a mechanistic representation of tumor evolution, from specific mutations to tumor size evolution. It additionally includes tumor shrinkage in response to administration of the first generation epidermal growth factor receptor tyrosine kinase inhibitor gefitinib (Lynch et al, 2004; Paez et al, 2004). We calibrated the ISELA model in order to predict tumor size evolution.

We describe here the two-step calibration that we applied to the ISELA model in order to ensure the model was reproducing the EGFR+ LUAD cell growth observed experimentally and described in literature.

### 3.2 Calibration protocol

#### 3.2.1 Parameterization of parameters

Within the ISELA model, the effect of mutations on the EGFR pathway was retrieved from the scientific literature and implemented as such, using parameterization. Here we present a compiled information on the main mutations that were parametrized prior to the calibration process:

- Kirsten rat sarcoma viral oncogene homolog gene (KRAS) mutation, which decreases by 97-99% (rounded to 98%) the hydrolysis of guanosine triphosphate once KRAS is activated, keeping it under its active form (Hunter et al, 2015)
- EGFR, exon 19 deletion, which has 3 distinct effects: it leads to the constitutive activation of EGFR (Harrison et al, 2020), it alters EGFR affinity for adenosine triphosphate from approximately 5.0μM to 129μM (Carey et al, 2006), and it modifies the affinity of EGFR to gefitinib changing the inhibition constant from approximately 16.4 nM to an estimated 0.833 nM based on an IC 50 of 6nM (Yasuda et al, 2013), making it sensitive to gefitinib treatment (Lynch et al, 2004)
- Phosphatidylinositol-4,5-Bisphosphate 3-Kinase Catalytic Subunit Alpha gene (PIK3CA) mutation, leading to an estimated 2.7 fold increase of PI3K complex activity (Yamamoto et al, 2008) conferring a resistance to gefitinib (Re et al, 2019)

#### 3.2.2 Assessing the calibration data for the ISELA model

We reviewed the scientific literature to retrieve the relevant experiments and biological phenomena related to the tumor growth in EGFR+ LUAD patients and their TTP. Five biological phenomena were identified as relevant to represent them in the context of development of the ISELA model:

1. tumor cell proliferation
2. tumor cell death
3. neoangiogenesis
4. effect of the immune system on the tumor
5. treatment

Together, these phenomena are represented with 14 parameters (see Supplementary Table S1), for which we aimed to find a single value corresponding to a mean behavior within the population.

A structured review of the relevant papers, given the context of use of the ISELA model, was performed. Table 1 lists the papers that were used for the calibration of tumor growth. These tumor growth-related experimental data can be divided in two categories:

1. ***In vitro* tumor growth data** (Jagiella et al, 2016). In this experimental context, only tumor cell proliferation and death can be followed. The main assumption is that the tumor can be represented as a spherical shape and grows proportionally (Lewin et al, 2018). The two remaining biological processes related to tumor growth, namely neoangiogenesis and the effect of the immune system on the tumor, are assumed not to operate in such an *in vitro* context and are thus set to null in the simulation.
2. ***In vivo* tumor xenograft mouse model growth data** (Kang et al, 2018). In this experimental context, tumor cell proliferation and death can be followed, as well as neoangiogenesis. Immunocompromised mice were ectopically xenografted with patient-derived NSCLC tumors. We selected data for such patient-derived tumors, one carrying EGFR exon 19 deletion mutation only, thus sensitive to gefitinib, one having both the exon 19 EGFR deletion and a mutation in PIK3CA (a subunit of PI3K complex, activating the PI3K/AKT pathway), thus resistant to gefitinib. Tumor size was measured once or twice per week using a vernier caliper. The effect of the immune system on the tumor is limited (due to immunosuppression), but not null. For the data we focus on mice treated with gefitinib at 25mg/kg or injected with 5 mM citrate buffer as a placebo, by oral gavage.

**Table 1.** References used for the calibration of tumor growth, focusing on implemented constraints on tumor size. § rim size = distance between tumor core and tumor surface

| Author (year) | Study type | Biological processes |
| --- | --- | --- |
| (Jagiella et al, 2016) | *in vitro* & *ex vivo* | Tumor spheroid growth curves, Fraction of proliferative cells at maximal radius |
| (Ekert et al, 2014) | *ex vivo* | Evolution of the proportion of proliferative cells within the tumor |
| (Freyer, 1988) | *in vitro* | Estimation of the viable rim size<sup>§</sup> of the tumor |
| (Kang et al, 2018) | *in vitro* | Tumor volume evaluation |

The experimental data used here present the following advantages:

- They are based on cells extracted from EGFR+ LUAD human tumors.
- Tumor growth is followed either without any treatment with gefitinib which matches the context of use of the ISELA model.
- The type of EGFR+ mutation of the tumor is specified.

#### 3.2.3 Ensuring that the model adequately represents data

Numbers of living, proliferative and dead cells are internal variables of the ISELA model. From these internal variables, the tumor volume and tumor radius are direct outputs of the model:

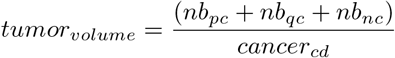

and

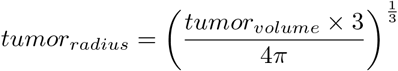

with *nb*_*pc*_ the number of proliferating cells, *nb*_*qc*_ the number of quiescent cells, *nb*_*nc*_ the number of necrotic cells, *cancer*_*cd*_ the density of cancer cells. For the sake of simplicity, *cancer*_*cd*_ is assumed to be constant within the tumor and over time and equals 2.8*e*^8^ cells/cm^3^ (Grassberger et al, 2019). Furthermore, tumors are assumed to be perfectly spherical. This hypothesis seems reasonable since the majority of tumors are spherical and approximately 20% are considered non-spherical (James et al, 1999). However, these assumptions may have an impact on the range of parameter values.

Consequently, under the two aforementioned assumptions, there is a direct match between the nature of the data reported in the selected papers, and the nature of the model outputs.

#### 3.2.4 Definition of the objective of the calibration steps

As mentioned previously, the goal of the calibration for the ISELA model was to reproduce the EGFR+ LUAD cell growth described in literature and observed experimentally.

The experimental data found in literature to characterize EGFR+ LUAD tumor growth are either obtained from *in vitro* spheroids, or from xenografted mice, reporting either tumor radius or tumor volume. We therefore decided to split the ISELA calibration into two sequential steps:

- Step 1: Reproduce the tumor radius evolution and the proportion of viable cells of *in vitro* spheroids. Step 1 addresses *in vitro* scale; it focuses on tumor growth in the absence of extracellular signals related to neoangiogenesis, immune system or treatment
- Step 2: Reproduce the tumor volume evolution of human EGFR+ LUAD tumor cells transplanted into immunocompromised mice. Step 2 addresses *in vivo* scale; it focuses on tumor growth in the presence of neoangiogenesis, immune system or treatment

While step 1 calibration is performed *in vitro*, step 2 is performed in mice. As a consequence calibration steps have different grain and scope. Another known consequence is that some of the reactions are expected to happen qualitatively at a different rate, for instance, doubling time of tumor cells are expected to be faster *in vitro*, than in mice, itself faster than in human ((Shimkin and Polissar, 1955), (Merk et al, 2009), (Arai et al, 1994), (Freyer, 1988)). We therefore calculated part of this experimental variability using allometry theory. Allometry describes how the characteristics of living creatures such as morphological and physiological traits change with their size. Kleiber’s law states that the metabolic rate scales to the 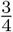 power of the animal’s mass for a wide range of species. Scaling laws are since then most often described with power law relationships *Z* = *a × M*^*b*^ with *Z* the studied characteristic, *M* the organism mass, and *a* and *b* parameters called allometric coefficient and allometric exponent respectively (Pérez-García et al, 2020; Smil, 2000), with the assumption that the allometric assumption relies on the physical dimension of the parameter ((Morgado et al, 1990), (Dawson, 2014), (West et al, 1997)). Allometric scaling thus provides a way to link parameters of individuals even when they are of different species. We decided to consider allometry in our model since this framework allows us to compare species on the basis of their body weight. For each parameter concerned by allometry, we calibrated the allometric coefficient *a* for a specific weight and this enabled us to deduce, using the power law relationships, the value of this parameter for different weight, having the allometric exponent *b* determined by the parameter physical unit. This allows one to maximize the use of the information obtained on one species on the other with the aim to have the model faithfully reproducing both, instead of doing independent calibration of the parameters in both species. In this study we used a reference body weight of 2.63g for *in vitro* experiments (West, 2002) and 23g for a mouse weight (Vellers et al, 2017).

#### 3.2.5 Automation of parameter value optimisation

Optimization of parameter values was performed automatically using the CMA-ES algorithm as described in the methods section.

### 3.3 Results

#### 3.3.1 Step 1: Reproduction of tumor growth (*in vitro* spheroid data)

The evolution of tumor radius prior and after calibration was compared to the experimental data used for calibration (see Figure 5). Figure 5 aims to provide a view on how the calibration outputs are at the same time:

- based on literature data knowledge, through the prior parameterization of the model, and could be able to provide acceptable results based on these considerations only
- improved by the model calibration process we developed here

**Fig. 5.**
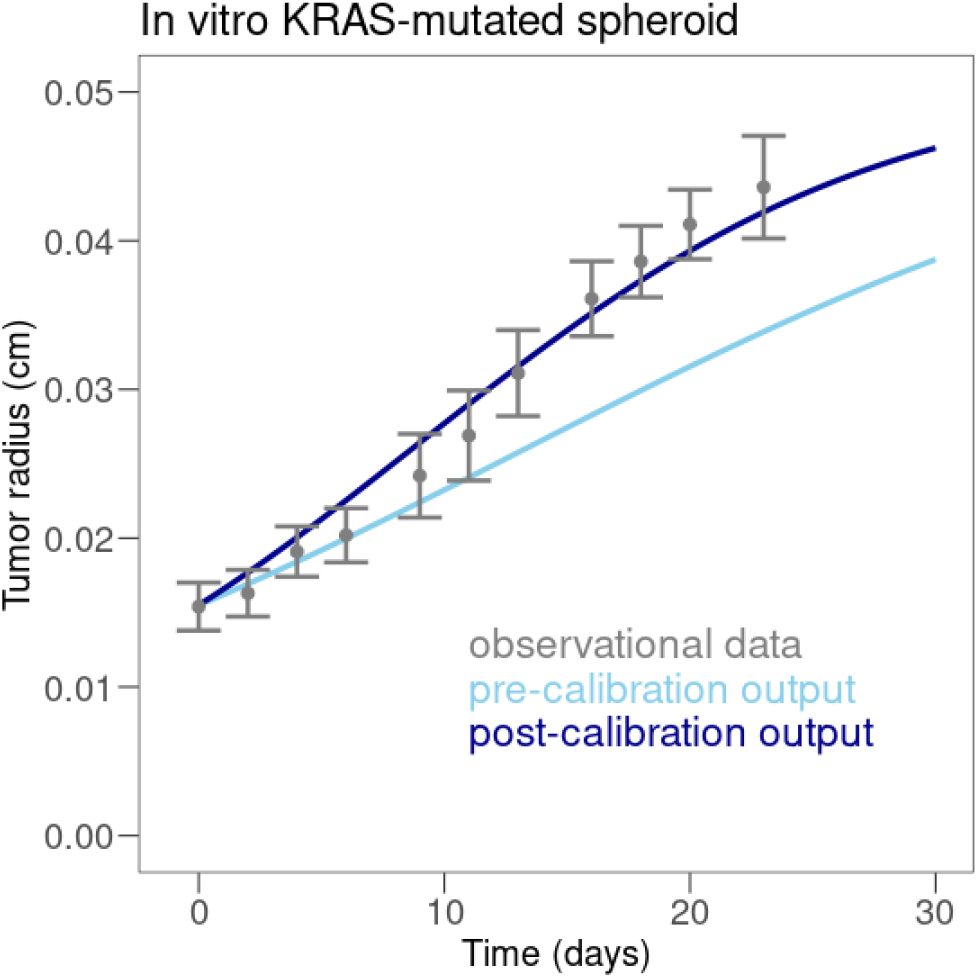
Efficacy of calibration of tumor radius on *in vitro* spheroids carrying KRAS mutation. Experimental data are shown in gray (error bars represent 2 standard deviations, experimental data are reported by Jagiella et al (2016), simulated data prior to calibration in light blue and simulated data after calibration in dark blue.

The experimental data showed an increase of tumor radius of approximately 12.5μm/day over the first 20 days, to reach 400μm, which then started to plateau. Pre-calibration of the model showed an increase of approximately 8μm/day. After calibration, as described in the calibration protocol section, the model showed an increase of approximately 12.5μm/day and the tumor radius stayed approximately within two standard deviations of the experimental data.

Overall, the calibration showed good agreement with experimental data post calibration, reproducing the expected growth of *in vitro* KRAS-mutated spheroids, at least for the first 30 days of the experiment.

#### 3.3.2 Step 2: Reproduction of tumor growth (*in vivo* xenograft growth data)

Evolution of tumor volume prior and after calibration was compared to the experimental data used for calibration for each of the four scenarios: EGFR mutation with or without gefitinib, EGFR and PIK3CA mutation with or without gefitinib (see Figure 6). In EGFR tumors, treated with placebo, the experimental data showed an increase of tumor volume of approximately 20mm^3^/day over the first 20 days (Figure 6, A). Under gefitinib in tumors with EGFR mutation only, volume decreased in size of roughly 16mm^3^/day over the first 10 days, before plateauing (Figure 6, B). In tumors with both EGFR and PIK3CA mutations, treatment with gefitinib showed no effect compared to placebo with tumor volume increasing roughly by 7.5mm^3^/day over the first 20 days (Figure 6, C and D).

**Fig. 6.**
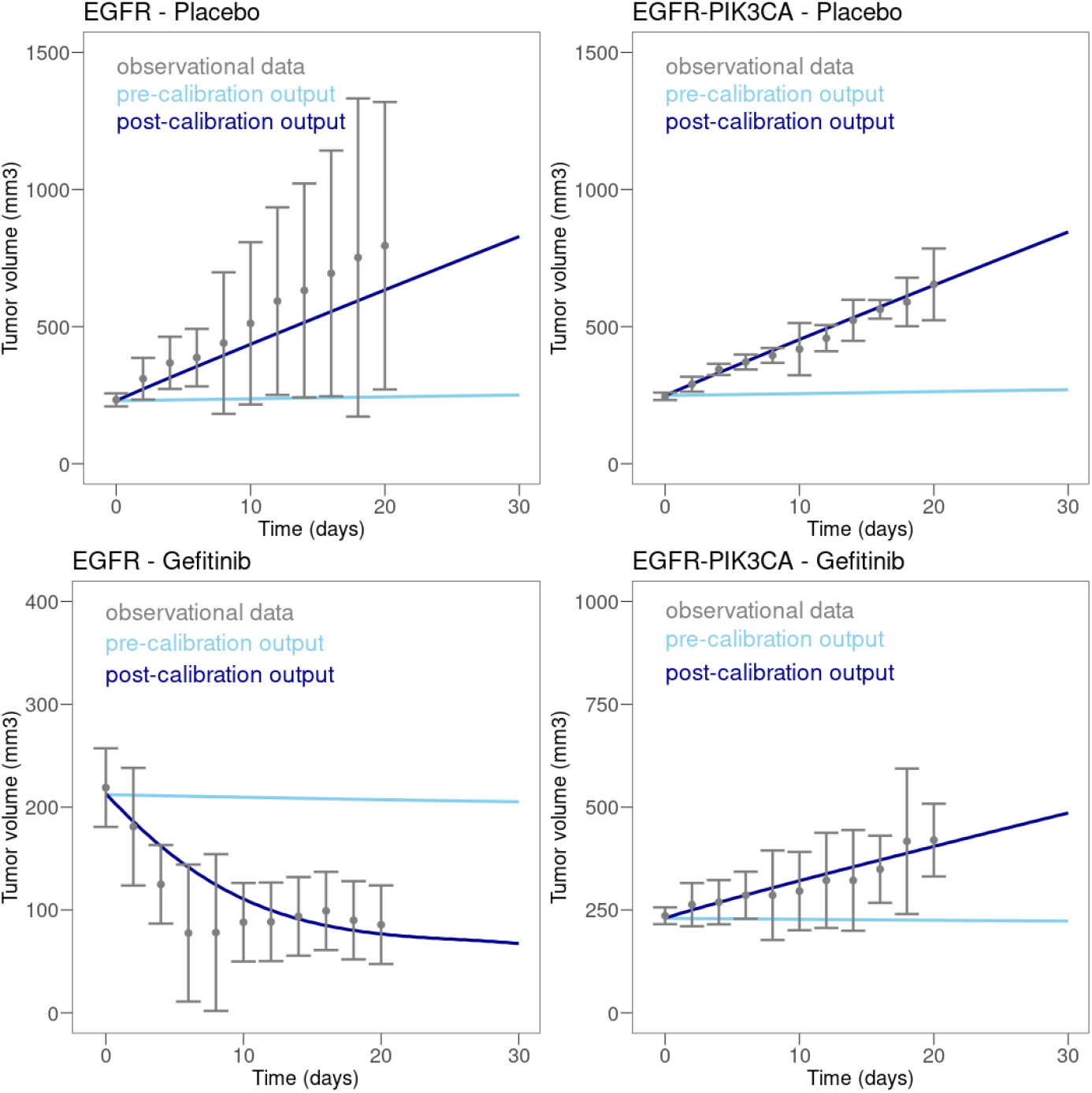
Efficacy of calibration of tumor volume with xenograft data. On each plot, the experimental data are shown in gray (Error bars on experimental data correspond to 2 standard deviations of the data reported in Kang et al (2018)), simulated data prior to calibration in light blue and simulated data after calibration in dark blue. The type of mutations carried by the tumor is indicated on top of each plot.

Overall, this calibration step successfully reproduced the growth of two xenografted tumors carrying EGFR exon 19 deletion, and one having in addition a mutation in PIK3CA, as reported in Kang et al (2018). The simulation could reproduce not only tumor evolution with placebo but also the differential effect of gefitinib depending on the tumor genotype.

The calibration thus succeeded in allowing the model to faithfully reproduce observed experimental data and fulfill the associated computational constraints. The changes in parameter values are detailed in the supplementary material.

## 4 Discussion

In mechanistic modeling, calibration is often seen as taking the risk of having a compensation of the model error by errors on the parameter values. We present here a calibration procedure improving model adequacy to both the knowledge and data extracted from literature. A perspective about knowledge-based models emphasizes the need to formalize model development and verification aspects and highlights optimal structural granularity and parameter estimation as critical aspects (Ribba et al, 2017). Calibration could also be necessary to estimate parameters value which can hardly be experimentally measured (*e*.*g*. R0 used in epidemiology (Delamater et al, 2019)). Calibration methodology, however, is often insufficiently discussed. If calibration is described, no standardized or generalized method is employed. Frequently, identifying the appropriate data set for calibration is a major challenge. Only a few papers could be identified with a detailed description of the calibration process. For instance, calibration was performed in a mathematical model of blood eosinophil dynamics against *in vitro* and *in vivo* data (Karelina et al, 2016). Willis et al (2021) have used parameter regression methods to calibrate an SIRD model for COVID-19. For a mechanistic model of oral poliovirus-related vaccine virus transmission risk, optimal model parameters were identified with Latin hypercube sampling and a pseudo-likelihood function to minimize the discrepancy between simulated and observed data (Famulare et al, 2021).

The methodology we presented here could maximize the use of heterogeneous data (in terms of scale, nature of measurement, qualitative or quantitative nature), to calibrate multiscale mechanistic model to better represent expected biological behavior observed at distinct scales.

From a theoretical perspective, the main limitations of such an approach are the availability of relevant knowledge and consistent data to efficiently constrain the model. In particular, when the biological constraints from literature are in disagreement, it may indicate that the respective phenomena are not sufficiently characterized in the literature. A documented removal of constraints or changes in the model structure may be viable options. Also, some phenomena may not be sufficiently documented in the literature, leading to a lack of computational constraints. In such a case, drawing some documented simplifying assumptions, or exploring alternative explanatory hypotheses, may be critical in model development and calibration. More generally, developing a more systematic way of dealing with incompatible constraints is an interesting avenue for future works.

The calibration strategy proposed in this paper can be compared to the use of machine learning (ML) in model development. ML model development is more standardized and includes training, testing and validation of a model. A training dataset is used initially to fit the model parameters. The model is then optimized on a training dataset with methods such as gradient descent to adjust model parameters. The fitted model is finally validated with a validation dataset where the model’s predictions are compared with the responses for the observations. In ML, similar to mechanistic modeling, lack of data is often the main limitation (Roh et al, 2021). Especially in healthcare, data-driven ML is limited due to bias and data access due to privacy concerns, data silos and heterogeneous data formats (Rieke et al, 2020).

From a technical perspective, there are two additional aspects that need to be considered for mechanistic modeling: the computational cost and the global convergence verification of the process. Calibration of many parameters requires significant computational power and a significant amount of available data. Therefore, prior to the calibration presented here, a parsimonious approach to model building with reducing the number parameters to calibrate (*e*.*g*. through simplifying assumptions, or sensitivity analysis to identify parameters affecting the outcome(s) of interest the most) is a key step. More fundamentally, the problem of parameter identifiability has been left unaddressed as well as the uncertainty quantification of the parameters after calibration. Both aspects will be the focus of future work. Also, a thorough exploration of calibration algorithm settings is necessary to ensure that the calibration converges to a unique set of values and that this convergence corresponds to a global maximum of the cost-function. Otherwise, if left unverified, the process may not have converged at all, or will have converged to a local maximum, leading to a suboptimal performance.

The stepwise process as presented here allows the calibration of increasingly larger models, starting from a relatively basic one to a more complex one. Those calibration protocols lead to the production of independent reusable subparts of models as well as complex models as a whole. These models can be reused in other contexts as a starting point to build calibrated mechanistic models adaptable to the available data and questions of interest. The method naturally produces calibrated portions of larger models, and these submodels can be used in other contexts.

We have applied this calibration strategy to a mechanistic multiscale KBM of lung adenocarcinoma harboring specific EGFR mutations, and focused on the calibration of tumor growth parameters based on *in vitro* experiments and xenografted mice. This led to the successful calibration of the model with these experimental settings as the calibrated model showed an adequate fit with the experimental data (see Figure 5 and 6). The model is currently under further development, in order to be able to represent measurements observed in humans. The ISELA model will likely require additional calibration steps focusing on specific parameters, *e*.*g*. to represent the human immune system. This would also imply calibrating not just the average behavior amongst the population, but also the inter-patient variability, to generate a Virtual Population matching the characteristics of a real population. To formally assess the performance of the model and its results’ generalizability, and after the calibration process, validation of the model will be performed with data that were neither used for building the model nor calibration, *e*.*g*. real world data or clinical trial data.

After the calibration process and an additional validation procedure on an independent dataset, an additional application of a mechanistic model would be the generation of independent virtual populations to answer clinical questions *in silico*.

## Acknowledgement

We would like to thank Janssen-Cilag France for supporting the project used as an example application in this article. We thank M. Margreiter for his valuable participation in the early phases of this project.

## Supplementary data

**Table S1.** Summary of parameters to calibrate, the phenomena they belong to and their values evolution over the course of the calibration process.

| Parameter description | Phenomena related to | Value prior to calibration (unit) | Value after calibration (unit) |
| --- | --- | --- | --- |
| Maximal depth of the tumor at which cells can survive without neoangiogenesis | tumor cell proliferation, tumor cell death | 104 ($\mu\text{m}$) | 111.8 ($\mu\text{m}$) |
| Ratio of living cells over proliferating ones with the tumor | tumor cell proliferation | 5 (unitless) | 4.52 (unitless) |
| Growth rate of the tumor | tumor cell proliferation | 0.07 ($\text{day}^{-1}$) | 0.0751 ($\text{day}^{-1}$) |
| Transduction of EGFR signaling into growth signaling, first parameter: equivalent to the signal required to reach half the maximal growth | tumor cell proliferation | 0.6 (unitless) | 0.416 (unitless) |
| Transduction of EGFR signaling into growth signaling, second parameter: equivalent to the sensitivity of tumor growth to change in EGFR signal | tumor cell proliferation | 10 (unitless) | 2.95 (unitless) |
| Death rate of the hypoxic tumor due to necrosis | tumor cell death | 1 ($\text{day}^{-1}$) | 0.435 ($\text{day}^{-1}$) |
| Transduction of EGFR signaling into cell survival signaling, first parameter: equivalent to the signal required to reach half the maximal growth | tumor cell death | 0.07 (unitless) | 0.0439 (unitless) |
| Transduction of EGFR signaling into cell survival signaling, second parameter: equivalent to the sensitivity of the cell survival to change in EGFR signal | tumor cell death | 10 (unitless) | 26.3 (unitless) |
| Maximal cell death inhibition the EGFR cell survival signaling can provide | tumor cell death | 0.8 (unitless) | 0.743 (unitless) |
| Number of cells neoangiogenesis can supply within the tumor | neoangiogenesis | 7.15e9 (cells) | 2.77e11 (cells) |
| Immune system cell killing, first parameter: maximal speed of tumor cell killing by the immune system | effect of the immune system on the tumor | 3.40e7 ($\text{cells.day}^{-1}$) | 5.94e9 ($\text{cells.day}^{-1}$) |
| Immune system cell killing, second parameter: affinity tumor cell killing by the immune system | effect of the immune system on the tumor | 1e9 (cells) | 9.89e10 (cells) |
| Immune system cell killing, third parameter: maximal number of cells the immune system can have access to | effect of the immune system on the tumor | 1e7 (cells) | 2.46e9 (cells) |
| Fraction of unbound gefitinib drug within the tumor | treatment | 1 (unitless) | 0.401 (unitless) |

## References

Allen R, Rieger T, Musante C (2016) Efficient generation and selection of virtual populations in quantitative systems pharmacology models. CPT: Pharmacometrics & Systems Pharmacology 5(3):140–146. https://doi.org/10.1002/psp4.12063

Arai T, Kuroishi T, Saito Y, et al (1994) Tumor doubling time and prognosis in lung cancer patients: evaluation from chest films and clinical follow-up study. japanese lung cancer screening research group. Jpn J Clin Oncol 24(4):199–204. https://doi.org/10.1093/oxfordjournals.jjco.a039706

Back T, Hammel U, Schwefel HP (1997) Evolutionary computation: Comments on the history and current state. IEEE transactions on Evolutionary Computation 1(1):3–17. https://doi.org/10.1109/4235.585888

Barber S, Voss J, Webster M (2015) The rate of convergence for approximate bayesian computation. Electronic Journal of Statistics 9(1):80–105. https://doi.org/10.1214/15-ejs988

Bhattacharya P, Viceconti M (2017) Multiscale modeling methods in biomechanics. Wiley Interdisciplinary Reviews: Systems Biology and Medicine 9(3):e1375. https://doi.org/10.1002/wsbm.1375

Bidot C, Monod H, Taupin ML (2018) A quick guide to multisensi, an r package for multivariate sensitivity analyses

Boissel JP, Auffray C, Noble D, et al (2015) Bridging systems medicine and patient needs. CPT: Pharmacometrics & Systems Pharmacology 4(3):135– 145. https://doi.org/10.1002/psp4.26

Brown KS, Sethna JP (2003) Statistical mechanical approaches to models with many poorly known parameters. Physical review E 68(2):021,904. https://doi.org/10.1103/physreve.68.021904

Brown LV, Gaffney EA, Wagg J, et al (2018) Applications of mechanistic modelling to clinical and experimental immunology: an emerging technology to accelerate immunotherapeutic discovery and development. Clinical and Experimental Immunology 193(3):284–292. https://doi.org/10.1111/cei.13182

Buil-Bruna N, López-Picazo JM, Martín-Algarra S, et al (2015) Bringing model-based prediction to oncology clinical practice: A review of pharmacometrics principles and applications. The Oncologist 21(2):220–232. https://doi.org/10.1634/theoncologist.2015-0322

Carey KD, Garton AJ, Romero MS, et al (2006) Kinetic analysis of epidermal growth factor receptor somatic mutant proteins shows increased sensitivity to the epidermal growth factor receptor tyrosine kinase inhibitor, erlotinib. Cancer Research 66(16):8163–8171. https://doi.org/10.1158/0008-5472.can-06-0453

Dawson T (2014) Allometric relations and scaling laws for the cardiovascular system of mammals. Systems 2(2):168–185. https://doi.org/10.3390/systems2020168, URL https://doi.org/10.3390/systems2020168,

Delamater PL, Street EJ, Leslie TF, et al (2019) Complexity of the basic reproduction number r0. Emerging Infectious Diseases 25(1):1–4. https://doi.org/10.3201/eid2501.171901, URL https://doi.org/10.3201/eid2501.171901

Delyon B, Lavielle M, Moulines E (1999) Convergence of a stochastic approximation version of the em algorithm. Annals of statistics pp 94–128. https://doi.org/10.1214/aos/1018031103

Duchesne R, Guillemin A, Crauste F, et al (2019) Calibration, selection and identifiability analysis of a mathematical model of the in vitro erythropoiesis in normal and perturbed contexts. In silico biology 13(1-2):55–69. https://doi.org/10.3233/isb-190471

Eissing T (2011) A computational systems biology software platform for multiscale modeling and simulation: Integrating whole-body physiology, disease biology, and molecular reaction networks. Frontiers in Physiology 2. https://doi.org/10.3389/fphys.2011.00004

Ekert JE, Johnson K, Strake B, et al (2014) Three-dimensional lung tumor microenvironment modulates therapeutic compound responsiveness in vitro – implication for drug development. PLoS ONE 9(3):e92.248. https://doi.org/10.1371/journal.pone.0092248

ESMO (2019) Non-small-cell lung cancer (NSCLC). ESMO Patient Guide Series -ESMO Clinical Practice Guidelines URL https://www.esmo.org/for-patients/patient-guides/non-small-cell-lung-cancer

Famulare M, Wong W, Haque R, et al (2021) Multiscale model for forecasting sabin 2 vaccine virus household and community transmission. PLOS Computational Biology 17(12):e1009.690. https://doi.org/10.1371/journal.pcbi.1009690

Freyer JP (1988) Role of necrosis in regulating the growth saturation of multicellular spheroids. Cancer Res 48(9):2432–2439. URL https://aacrjournals.org/cancerres/article-pdf/48/9/2432/2434811/cr0480092432.pdf

Galtier MN, Wainrib G (2013) A biological gradient descent for prediction through a combination of stdp and homeostatic plasticity. Neural computation 25(11):2815–2832. https://doi.org/10.1162/neco_a_00512

Grassberger C, McClatchy D, Geng C, et al (2019) Patient-specific tumor growth trajectories determine persistent and resistant cancer cell populations during treatment with targeted therapies. Cancer research 79(14):3776–3788. https://doi.org/10.1158/0008-5472.CAN-18-3652

Hansen N, Kern S (2004) Evaluating the cma evolution strategy on multimodal test functions. In: Yao X, Burke EK, Lozano JA, et al (eds) Parallel Problem Solving from Nature - PPSN VIII. Springer Berlin Heidelberg, Berlin, Heidelberg, pp 282–291, https://doi.org/10.1007/978-3-540-30217-9_29

Hansen N, Müller SD, Koumoutsakos P (2003) Reducing the time complexity of the derandomized evolution strategy with covariance matrix adaptation (CMA-ES). Evolutionary Computation 11(1):1–18. https://doi.org/10.1162/106365603321828970

Hansen N, Akimoto Y, Baudis P (2019) CMA-ES/pycma on Github https://doi.org/10.5281/zenodo.2559634

Harrison PT, Vyse S, Huang PH (2020) Rare epidermal growth factor receptor (EGFR) mutations in non-small cell lung cancer. Seminars in Cancer Biology 61:167–179. https://doi.org/10.1016/j.semcancer.2019.09.015

Horstemeyer MF (2009) Multiscale modeling: A review pp 87–135. https://doi.org/10.1007/978-90-481-2687-3_4

Hunter JC, Manandhar A, Carrasco MA, et al (2015) Biochemical and structural analysis of common cancer-associated KRAS mutations. Molecular Cancer Research 13(9):1325–1335. https://doi.org/10.1158/1541-7786.mcr-15-0203

Jagiella N, Müller B, Müller M, et al (2016) Inferring growth control mechanisms in growing multi-cellular spheroids of NSCLC cells from spatialtemporal image data. PLOS Computational Biology 12(2):e1004.412. https://doi.org/10.1371/journal.pcbi.1004412

James K, Eisenhauer E, Christian M, et al (1999) Measuring response in solid tumors: Unidimensional versus bidimensional measurement. JNCI Journal of the National Cancer Institute 91(6):523–528. https://doi.org/10.1093/jnci/91.6.523, URL https://doi.org/10.1093/jnci/91.6.523

Jones D, Floudas C, Pardalos P (2001) Encyclopedia of optimization. DIRECT global optimization pp 725–735. https://doi.org/10.1007/978-0-387-74759-0_128

Jdrzejewski-Szmek Z, Abrahao KP, Jdrzejewska-Szmek J, et al (2018) Parameter optimization using covariance matrix adaptation—evolutionary strategy (CMA-ES), an approach to investigate differences in channel properties between neuron subtypes. Frontiers in Neuroinformatics 12. https://doi.org/10.3389/fninf.2018.00047

Kang HN, Choi JW, Shim HS, et al (2018) Establishment of a platform of nonsmall-cell lung cancer patient-derived xenografts with clinical and genomic annotation. Lung Cancer 124:168–178. https://doi.org/10.1016/j.lungcan.2018.08.008

Karelina T, Voronova V, Demin O, et al (2016) A mathematical modeling approach to understanding the effect of anti-interleukin therapy on eosinophils. CPT: Pharmacometrics & Systems Pharmacology 5(11):608– 616. https://doi.org/10.1002/psp4.12129

Kirkpatrick S, Gelatt Jr CD, Vecchi MP (1983) Optimization by simulated annealing. science 220(4598):671–680. https://doi.org/10.1126/science.220.4598.671

Klipp E, Liebermeister W, Wierling C, et al (2016) Systems biology: a textbook. John Wiley & Sons

Lewin TD, Maini PK, Moros EG, et al (2018) The evolution of tumour composition during fractionated radiotherapy: Implications for outcome. Bulletin of Mathematical Biology 80(5):1207–1235. https://doi.org/10.1007/s11538-018-0391-9

Liepe J, Filippi S, Komorowski M, et al (2013) Maximizing the information content of experiments in systems biology. PLoS computational biology 9(1):e1002.888. https://doi.org/10.1371/journal.pcbi.1002888

Liu JS, Chen R (1998) Sequential monte carlo methods for dynamic systems. Journal of the American statistical association 93(443):1032–1044. https://doi.org/10.1080/01621459.1998.10473765

Lynch TJ, Bell DW, Sordella R, et al (2004) Activating mutations in the epidermal growth factor receptor underlying responsiveness of non-small-cell lung cancer to gefitinib. N Engl J Med 350(21):2129–2139. https://doi.org/10.1056/NEJMoa040938

Merk J, Rolff J, Becker M, et al (2009) Patient-derived xenografts of non-small-cell lung cancer: a pre-clinical model to evaluate adjuvant chemotherapy? European Journal of Cardio-Thoracic Surgery 36(3):454–459. https://doi.org/10.1016/j.ejcts.2009.03.054

Miao H, Xia X, Perelson AS, et al (2011) On identifiability of nonlinear ode models and applications in viral dynamics. SIAM review 53(1):3–39. https://doi.org/10.1137/090757009

Moles CG, Mendes P, Banga JR (2003) Parameter estimation in biochemical pathways: a comparison of global optimization methods. Genome research 13(11):2467–2474. https://doi.org/10.1101/gr.1262503

Monod H, Bouvier A, Kobilinsky A (2012) A quick guide to planor, an r package for the automatic generation of regular factorial designs. Tech. rep., Citeseer

Morgado E, Ocqueteau C, Cury M, et al (1990) Three-dimensional morphometry of mammalian cells. II. areas, volumes, and area-volume ratios. Arch Biol Med Exp (Santiago) 23(1):21–27

Morrison TM, Hariharan P, Funkhouser CM, et al (2019) Assessing computational model credibility using a risk-based framework: Application to hemolysis in centrifugal blood pumps. ASAIO Journal 65(4):349–360. https://doi.org/10.1097/mat.0000000000000996

Musante C, Ramanujan S, Schmidt B, et al (2016) Quantitative systems pharmacology: A case for disease models. Clinical Pharmacology & Therapeutics 101(1):24–27. https://doi.org/10.1002/cpt.528

Nelder JA, Mead R (1965) A simplex method for function minimization. The computer journal 7(4):308–313. https://doi.org/10.1093/comjnl/7.4.308

Paez JG, Jänne PA, Lee JC, et al (2004) EGFR mutations in lung cancer: Correlation with clinical response to gefitinib therapy. Science 304(5676):1497–1500. https://doi.org/10.1126/science.1099314

Pant S, Lombardi D (2015) An information-theoretic approach to assess practical identifiability of parametric dynamical systems. Mathematical biosciences 268:66–79. https://doi.org/10.1016/j.mbs.2015.08.005

Pérez-García VM, Calvo GF, Bosque JJ, et al (2020) Universal scaling laws rule explosive growth in human cancers. Nature physics 16(12):1232–1237. https://doi.org/10.1038/s41567-020-0978-6

Re MD, Crucitta S, Gianfilippo G, et al (2019) Understanding the mechanisms of resistance in EGFR-positive NSCLC: From tissue to liquid biopsy to guide treatment strategy. International Journal of Molecular Sciences 20(16):3951. https://doi.org/10.3390/ijms20163951

Renardy M, Hult C, Evans S, et al (2019) Global sensitivity analysis of biological multiscale models. Current opinion in biomedical engineering 11:109–116. https://doi.org/10.1016/j.cobme.2019.09.012

Ribba B, Grimm H, Agoram B, et al (2017) Methodologies for quantitative systems pharmacology (QSP) models: Design and estimation. CPT: Pharmacometrics & Systems Pharmacology 6(8):496–498. https://doi.org/10.1002/psp4.12206

Rieke N, Hancox J, Li W, et al (2020) The future of digital health with federated learning. npj Digital Medicine 3(1). https://doi.org/10.1038/s41746-020-00323-1

Roh Y, Heo G, Whang SE (2021) A survey on data collection for machine learning: A big data - AI integration perspective. IEEE Transactions on Knowledge and Data Engineering 33(4):1328–1347. https://doi.org/10.1109/tkde.2019.2946162

Rosenthal JS (1995) Minorization conditions and convergence rates for markov chain monte carlo. Journal of the American Statistical Association 90(430):558–566. https://doi.org/10.1080/01621459.1995.10476548

Ryan PIC, et al (2007) References to cma-es applications. Strategies 4527(467)

Shimkin MB, Polissar MJ (1955) Some quantitative observations on the induction and growth of primary pulmonary tumors in strain a mice receiving urethan. J Natl Cancer Inst 16(1):75–97. https://doi.org/10.1093/jnci/16.1.75

Smil V (2000) Laying down the law. Nature 403(6770):597–597. https://doi.org/10.1038/35001159

Tomasoni D, Paris A, Giampiccolo S, et al (2021) QSPcc reduces bottlenecks in computational model simulations. Communications Biology 4(1). https://doi.org/10.1038/s42003-021-02553-9

Toni T, Welch D, Strelkowa N, et al (2009) Approximate bayesian computation scheme for parameter inference and model selection in dynamical systems. Journal of the Royal Society Interface 6(31):187–202. https://doi.org/10.1098/rsif.2008.0172

Vellers HL, Letsinger AC, Walker NR, et al (2017) High fat high sugar diet reduces voluntary wheel running in mice independent of sex hormone involvement. Frontiers in physiology 8:628. https://doi.org/10.3389/fphys.2017.00628

West G (2002) Woodruff wh, brown jh. Allometric scaling of metabolic rate from molecules and mitochondria to cells and mammals Proc Natl Acad Sci USA 99:2473–2478. https://doi.org/10.1073/pnas.012579799

West GB, Brown JH, Enquist BJ (1997) A general model for the origin of allometric scaling laws in biology. Science 276(5309):122–126. https://doi.org/10.1126/science.276.5309.122, URL https://doi.org/10.1126/science.276.5309.122

Willis MJ, Wright A, Bramfitt V, et al (2021) COVID-19: Mechanistic model calibration subject to active and varying non-pharmaceutical interventions. Chemical Engineering Science 231:116,330. https://doi.org/10.1016/j.ces.2020.116330

Yamamoto H, Shigematsu H, Nomura M, et al (2008) PIK3ca mutations and copy number gains in human lung cancers. Cancer Research 68(17):6913– 6921. https://doi.org/10.1158/0008-5472.CAN-07-5084

Yasuda H, Park E, Yun CH, et al (2013) Structural, biochemical, and clinical characterization of epidermal growth factor receptor (EGFR) exon 20 insertion mutations in lung cancer. Science Translational Medicine 5(216). https://doi.org/10.1126/scitranslmed.3007205

Yugi K (2013) Dynamic kinetic modeling of mitochondrial energy metabolism pp 105–142. https://doi.org/10.1007/978-1-4614-6157-9_8

